# Limited value of Nanopore adaptive sampling in a long-read metagenomic profiling workflow of clinical sputum samples

**DOI:** 10.1101/2025.03.25.645176

**Authors:** Weizhen Xu, Janetta Top, Mattheus C. Viveen, Andrii Slyzkyi, Noud Hermans, Sarah van Erp, Dafna Eiloz, Richard Anthony, Kristin Kremer, Anita C. Schürch

## Abstract

Oxford Nanopore adaptive sampling (NAS) is a method by which the long-read sequencing flowcell accepts or rejects DNA molecules that are actively being sequenced based on their initial ∼500bp sequences, selectively increasing target data output. NAS promises up to 5-10x enrichment of target sequencing yield without additional sample preparation, but this optimal performance is dependent on ideal sample parameters which may be difficult to achieve under many real-world use-cases. We evaluated the use of NAS with the current R10.4.1 flowcell chemistry for profiling clinical sputum metagenomes, achieving at best 3.1× enrichment of bacterial sequence output due to the shorter read lengths (∼2.5kb) from the PCR amplification necessary to compensate for low DNA extraction yields. More critically, we encountered rapid pore loss during our runs that reduced total sequencing yield by an estimated 80%. We were unable to mitigate the pore loss despite extensive attempts to reduce contaminant carry-over, and we could not determine its cause but ruled out NAS and pore underloading as contributing factors. We conclude that the utility of NAS is often limited by the characteristics of the metagenomic sample studied, and that the factors contributing to pore loss need to be resolved before ONT sequencing can be reliably applied to long-read metagenomics.

## Introduction

Culture-based isolate sequencing followed by phenotypic testing of drug susceptibility is generally regarded to be the gold-standard practice for the epidemiological surveillance of infectious diseases and the associated spread of antimicrobial resistance and virulence genes. However, this process is slow and complex, especially for difficult-to-culture or slow-growing pathogens such as *Mycobacterium tuberculosis*. Furthermore, it relies on well-staffed centralised laboratory infrastructures that are more difficult to establish in lower-middle income country settings where the burden of infectious disease is the greatest(1–4). Direct culture-free metagenomic sequencing of clinical samples would avoid the aforementioned difficulties with microbial culture and offers a comprehensive genomic overview of multiple species present at the site of infection(1, 3–5), but is complicated by high levels of off-target host or commensal DNA (up to 99.99% in respiratory samples) that often hinder the detection of low-abundance species and genes of interest(1, 3, 4, 6, 7). Chemical host-depletion methods can improve metagenomic sensitivity and performance(3, 4, 6) but impose additional costs and workflow complexity that might be challenging in low-resource settings.

Nanopore adaptive sampling (NAS) is a methodology within Oxford Nanopore Technologies (ONT) sequencing that enriches for target sequences without additional wet lab preparation. NAS utilises the real-time sequencing capabilities of ONT to read the first 400-500bp of a DNA molecule, comparing it against a database of target or non-target genomic sequences and rejecting off-target molecules accordingly, preferentially sequencing target molecules at full length(8, 9). NAS enrichment performance is dependent on multiple sample parameters, in particular longer (>10kb) read length(8). While the ONT NAS best-practices guide(10) promises up to 5-10× target enrichment, occasionally achievable when higher-quality DNA samples can be obtained(8, 11, 12), enrichment performance was lower (1.8-4×) in many other NAS metagenomic studies(9, 13–20), likely reflective of real-world samples deviating from optimal parameters. These studies were done using the earlier generation R9 ONT flowcell chemistry in which NAS was first developed and optimised(9). We thus evaluated the use of the current generation of R10.4.1 flowcells in a multiplexed NAS-based workflow for the metagenomic and antimicrobial resistance (AMR) gene profiling of actual clinical sputum samples in a low-resource laboratory setting. While easily collectible in a minimally invasive manner, the higher host DNA composition and low extraction yields of sputum present a greater challenge for metagenomic profiling.

We encountered rapid pore loss in the ONT flowcells during this evaluation, severely limiting metagenomic performance. Pore degradation either from electric potential(21) or pore blockage(22) is a major factor affecting ONT sequencing yield and inter-run variability(8), and while accelerated pore loss is a commonly-cited cause of run failure on the ONT community forums (closed access to registered ONT customers, 2023-2025), its specific causes and solutions are not well-documented in scientific literature. We thus present our efforts to troubleshoot pore loss based on ONTs’ recommendations of improving pore occupancy and reducing sample contamination.

### Materials and methods Sputum collection

The sputum samples used in this study included residual and completely anonymized material from routine diagnostic protocols at University Medical Centre Utrecht (UMCU). Ethical approval for the use of the sputum samples for this study was obtained from the UMCU Biobank Research Ethics Committee (TCBio) with approval number: TCBio 25-063.

### Mock respiratory pathogen community

A mock community of respiratory pathogens was prepared from clinical isolates of *Pseudomonas aeruginosa* and *Klebsiella pneumoniae* from our hospital and type cultures of *Staphylococcus aureus* USA300 and *Streptococcus pneumoniae* ATCC49619. Glycerol stocks of these isolates were initially plated and incubated overnight on tryptone-soy-agar (TSA) (BD, Alphen aan den Rijn, The Netherlands) at 37°C, and in the case of *S. pneumoniae* at 37°C in a 5% CO_2_ atmosphere incubator. Colonies of the first three pathogens were inoculated into 10ml Luria Broth (LB) medium (Sigma Aldrich, Zwijndrecht, The Netherlands) and cultured overnight at 37°C, while *S. pneumoniae* was subcultured on four TSA plates at 37°C in a 5% CO_2_ atmosphere overnight and resuspended in 10ml LB medium. Bacterial load from these cultures was estimated based on OD_600_ absorbance (OD_600_ of 1.0 = 1×10^10^ cfu), pooled at equal culture densities and aliquoted into 50μl volumes of 3.1×10^8^ cfu total bacteria each. In the pilot run, ten-fold serial dilutions (3.1×10^7^-10^4^ cfu) of this mock community stock were added to each sample from two clinical sputa for subsequent spike-in experiments.

### High molecular weight DNA extraction from sputum

Approximately 200μl of each sputum sample was mixed with an equal volume of SPUTOLYSIN® reagent (Merck), resuspended with a Vortex Genie-2 laboratory vortex mixer (Scientific Industries) and incubated at 37°C with shaking at 800rpm for 15min to obtain homogenous suspensions. The samples were then centrifuged at 16,060g for 5min, after which the supernatant was discarded and the pellets resuspended in 300μl sterile 1× Dulbecco’s phosphate-buffered saline (DPBS, Capricorn Scientific) each.

Bacteria lysis was performed by adding an enzymatic mix of 5μl achromopeptidase (50000U/ml, Merck), 1.5μl lysostaphin (10mg/ml, Sigma), 20μl lysozyme (50mg/ml, Merck) and 6μl mutanolysin (2000U/ml, Sigma) to each sample, and incubating the samples at 37°C with shaking at 800rpm for 2h. Further mechanical lysis and DNA extraction was performed using the DNeasy UltraClean Microbial kit (QIAGEN). We adapted the mechanical lysis step for easier implementation in low-resource lab settings, adding 300µl of PowerBead solution from the DNeasy kit to each sample and vortexing at maximum speed with the same Vortex Genie-2 vortex mixer with a vortex adaptor for 24 1-1.5ml tubes(QIAGEN). DNA was subsequently extracted according to standard kit protocol and dsDNA concentrations were quantified on a Qubit™ 2 fluorometer using the Qubit™ high sensitivity quantification assay kit (ThermoFisher). High molecular weight DNA size was assessed in earlier extractions by the Genomic DNA ScreenTape assay on the TapeStation electrophoresis system (Agilent).

A negative control consisting of 300μl sterile 1× DPBS subjected to the same bacteria lysis treatment and DNA extraction procedure was included in each run.

### Initial library preparation and sequencing

For the first two ONT sequencing runs, multiplexed libraries were prepared with the Ligation Sequencing Kit V14 (SQK-LSK114) with PCR Expansion (EXP-PCA001) according to the ONT ligation sequencing of low input gDNA protocol (version LWP_9183_v114_revG_07Mar2023), using a PCR template input of up to 100ng depending on sample yield and 10× PCR extension cycles of 8min each, using the LongAMP Taq 2× Master Mix (NEB). Barcoded samples were quantified by Qubit™ fluorometer and combined into a equimolar pooled library, with the volume of the negative control sample matching that of the lowest concentration sample. ONT R10.4.1 flowcells (FLO-MIN114) with included light shields were used during sequencing for a duration of 72h. For the first pilot run, 150ng (∼75fmol) of pooled library was loaded, with no subsequent need for sample reloading or flowcell washing. For the second run, 125ng (∼64fmol) of library was loaded without the nuclease wash step for the first flowcell (2A), and with a nuclease wash/reload of 125ng library pool for the second (2B) at 24h.

Flowcells were run on a ONT GridION x5 Mk1 platform using the Dorado 7.3.11 high-accuracy basecalling model (400bps). The pilot run was performed using MinKNOW software v23.11.7, and the second run using MinKNOW v24.02.16. NAS was implemented through the MinKNOW interface to deplete host reads aligning to the human T2T-CHM13v2 genome reference (GCF_009914755.1). In the pilot run, channels 1-256 were designated as NAS channels and channels 257-512 as control non-NAS channels. All channels were used for NAS in the second run.

### Troubleshooting run library preparation and sequencing

We revised the earlier library preparation protocol to address potential contaminant carryover, flowcell underloading and to troubleshoot possible pore blockage resulting from NAS. Multiplexed libraries were prepared using the Rapid PCR Barcoding Kit 24 V14 (SQK-RPB114.24) according to ONT protocol (version RPB_9191_v114_revB_28Jun2023), using a PCR template input of up to 4ng DNA and 30× PCR extension cycles of 6min each. Samples were combined into an equimolar pool as in the pilot runs.

To troubleshoot if there were any effects of NAS on pore health, we loaded 400ng (∼200fmol) each of the same library pool onto four separate ONT R10.4.1 flowcells to maximize pore occupancy, with two flowcells running NAS on half the channels similar to the pilot run, and two control flowcells not running NAS. In the second set of replicates for the control/NAS runs, we added 0.25μl of RNase A (10μg/ml, QIAGEN, RP14) to the library pool for 15min at room temperature before loading the flowcell to troubleshoot possible RNA contamination degrading flowcell performance.

Flowcells were run for 72h with a nuclease wash-reload of 200ng of the library pool (∼100fmol) at 24h on the GridION platform using the same software versions as the second run (MinKNOW v23.11.7 and Dorado 7.3.11 high-accuracy basecalling).

### Bioinformatic analysis

We initially demultiplexed basecalled .fastq reads by barcode and read quality in the MinKNOW interface. Total flowcell data output was calculated from successfully barcoded reads with a minimum Q score of 9, inclusive of short (∼500bp) NAS-rejected reads. NAS-level read demultiplexing was performed using custom in-house scripts available at https://gitlab.com/weizhenxu/pas4amr-tools, with read output demultiplexed based on whether the read was obtained from a control or NAS channel (channel_demultiplex.py), or if it was rejected (“reject”) or accepted (“keep”) by NAS (unblock_demultiplex.py) based on the MinKNOW generated adaptive sampling report.

Reads were taxonomically classified using Kraken 2 (v2.1.3) (23) using the standard-16 database available at https://benlangmead.github.io/aws-indexes/k2 (last accessed 9 Jan 2024). A species was considered to detected in a sample if >=100 reads were assigned to it at the species (“S”) rank in the sample report. In samples where reads were assigned to multiple closely-related species in the same genus, identified species with <10% of the read count of the most abundant species in that genus were considered to be cross-mapped and removed in the final analysis.

Acquired genes and chromosomal point mutations linked to AMR were detected by ResFinder v4.5.0(24) using the ResFinder 2.02 and PointFinder 4.1.0 databases, with identity and alignment coverage thresholds of >95% each.

### Data availability

Raw sequencing data was processed using Hostile v2.0.0 with the human-t2t-hla-argos985 index (25) to remove human origin reads, and the processed .fastq files were uploaded and made available to the European Nucleotide Archive (ENA) project PRJEB86674.

## Results

### Reduced bacterial enrichment by NAS of PCR-barcoded libraries

DNA extraction yields from sputa ranged from 7.4 to 46.2 ng/μl and were often below the recommended concentration (<40ng/μl) for a ligation-based native barcoding library preparation workflow. We thus opted for the PCR barcoding expansion kit (EXP-PCA001)-based protocol to accommodate low-yield samples and to improve equimolar loading of multiplexed samples.

The aim of our pilot run was to assess the practical detection limits of pathogen genomes in actual clinical sputa with NAS and to accordingly determine the target read yields necessary for robust metagenomic profiling. We sequenced a mock community of four equally-pooled respiratory bacterial pathogens (Gram-negative: *Klebsiella pneumoniae* and *Pseudomonas aeruginosa*, Gram-positive: *Staphylococcus aureus* and *Streptococcus pneumoniae*) spiked at ten-fold dilutions at 3.1×10^7^-3.1×10^4^ cfu into two clinical sputa. This spike series, unspiked sputa and positive and negative controls were sequenced in multiplex on a single MinION flowcell divided in half into NAS channels (channel 1-256) and control channels (channel 257-512) to evaluate the relative enrichment benefit of NAS (**Table S1**). The total sequencing output (including short ∼500bp rejected reads by NAS) was 9.62M reads/13.7Gb passing quality thresholds (Q > 9) (**Table 1**, **Fig. S1A**). The PCR amplification resulted in shorter reads compared to the input template (previously determined to 15-30kb by TapeStation sizing) despite long 8min extension cycles, indicated by a ∼2.5kb modal peak corresponding to the non-rejected reads (**Fig. S1B).**

**Table 1.** Summary statistics of first two nanopore adaptive sampling (NAS) runs.

|  | Total | Pilot run |  | Run 2 (100% NAS run) |  |  |
| --- | --- | --- | --- | --- | --- | --- |
|  |  | Control channels | NAS channels | Total pool | Flowcell 2A | Flowcell 2B |
| **Total read count (M)** | **9.62** | 2.43 | 7.19 | 4.18 | 1.89 | 2.19 |
| **Total base output (Gb)** | **13.7** | 6.9 | 6.8 | 5.0 | 2.2 | 2.8 |
| **Human reads (M)** | **9.19** | 2.30 | 6.79 | 3.59 | 1.67 | 1.92 |
| **Bacterial reads (M)** | **0.52** | 0.13 | 0.39 | 0.49 | 0.22 | 0.27 |
| **% bacterial reads** | **5.4** | 5.2 | 5.5 | 11.9 | 11.6 | 12.4 |
| **On-cell target enrichment** |  | **3.1x** |  | **N.A. (no control channels)** |  |  |

The total sequencing output (including rejected reads) of the control and NAS channels were 6.9Gb/6.8Gb respectively. Similar to the previous metagenomic study by Marquet et al.(14), NAS had a minor effect on the proportional count of bacterial reads sequenced (5.5% NAS reads vs 5.2% control reads, or a 1.06× increase) – NAS enrichment was a result of increased read turnover in NAS channels due to the earlier rejection of host reads, resulting in 3.0× more total reads and 3.1× as more bacterial reads (**Table 1**), comparable to other PCR-based NAS metagenomic studies(14, 19, 26) but less than the 5-10× optimal enrichment suggested in the ONT best practices guide and observed in studies using longer non-PCR reads(8, 11, 12).

We analysed the taxonomic composition of the “keep” and “reject” fractions of the NAS channel output to determine NAS decision accuracy (**Table S2**). The overall accuracy of NAS was 94.9%, with similar proportions of falsely-accepted human reads (5.2%) and falsely-rejected bacterial reads (5.1%). Given the high proportion of host reads in the input (94.8% based on the control channels), non-rejected human reads accounted for 48% of the “keep” fraction. In agreement with Marquet et al(14)., NAS alone was insufficient to comprehensively remove human reads from the final sequencing output, necessitating the use of more stringent software host-removal tools(25) to avoid potential patient privacy issues.

Analysis of the sequencing output from the positive control consisting of the mock community alone revealed a disproportionate read yield for the two Gram-negative species accounting for >90% of the total reads (**Table S3**). This possibly reflects a bias in DNA extraction efficiency for Gram-positive and -negative species. Dilution of the mock community spike in the sputa resulted in ∼100× and ∼20× less reads by proportion relative to the undiluted positive control or 10^3^-10^4^ reads per mock community species in the samples with the highest 3.1×10^7^ cfu total spike, and <100 reads per million for most of the pathogenic species in the 3.1×10^5^ cfu or lower mock community spikes (**Fig. 1**). We tested if AMR genotypic profiles from previous in-house assemblies of the *K. pneumoniae* and *P. aeruginosa* mock community isolates could be reproduced from our NAS sequencing data (**Fig. 2**). Most of the expected AMR genes from the assemblies could be recovered in the positive control sample (*K. pneumoniae:* 16/17 acquired AMR genes and 4/4 AMR-linked chromosomal point mutations, *P. aeruginosa:* 10/11 acquired AMR genes), but only partial profiles were detected in 3.1×10^7^ cfu spiked sputum B (*K. pneumoniae:* 13/17 acquired genes and 2/4 chromosomal point mutations, *P. aeruginosa:* 5/11 acquired AMR genes), 1/17 *K. pneumoniae* acquired genes detected in the 3.1×10^7^ cfu spike of sputum A, and no genes detected at the lower spike concentrations/species read counts.

**Fig. 1.**
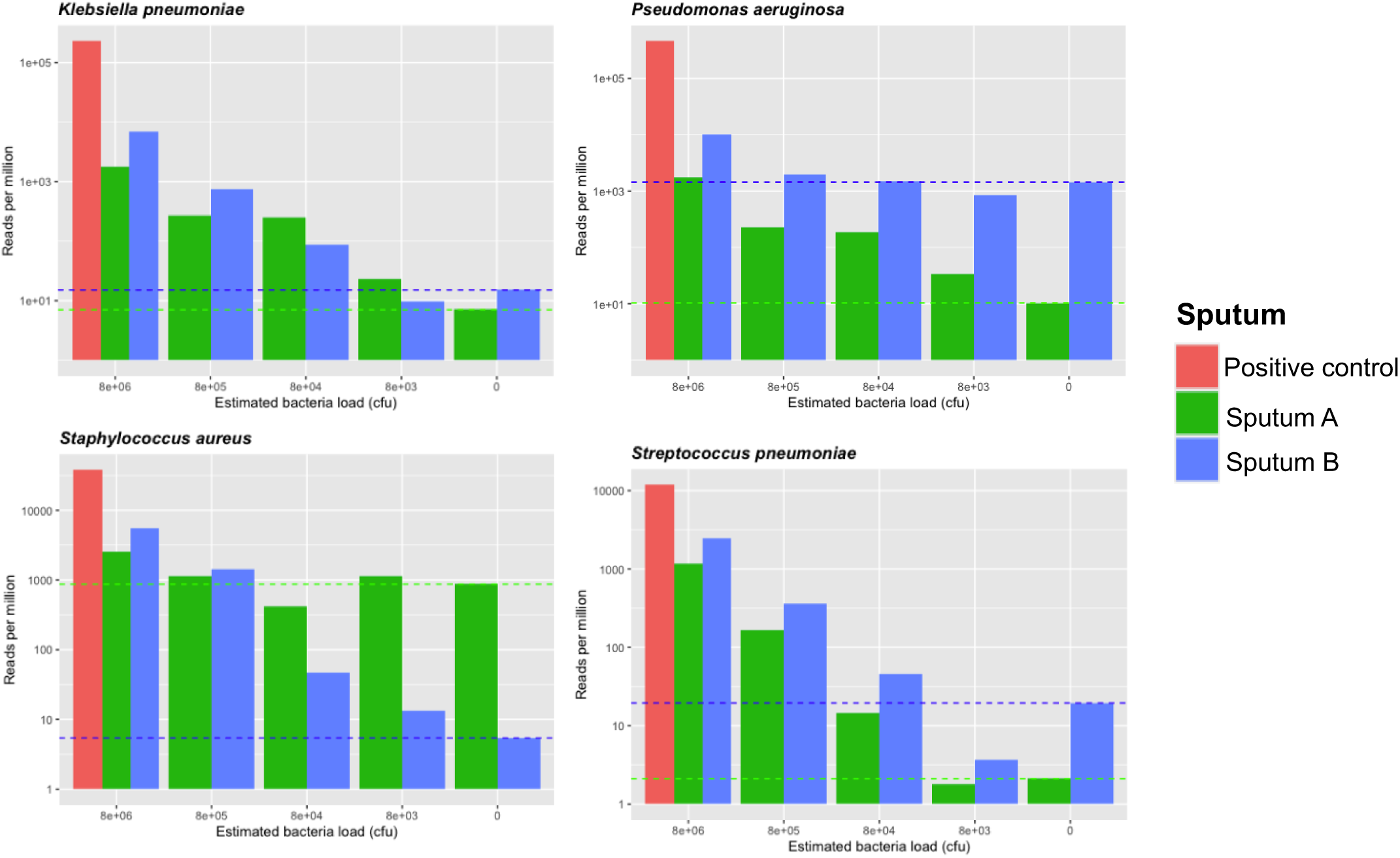
Detection of mock respiratory pathogen community reads in spiked sputa. Bars indicate reads from the mock community positive control without sputum (pink) and sputa A (green) and B (blue). Dotted lines represent baseline counts from DNA already present in unspiked sputa.

**Fig. 2.**
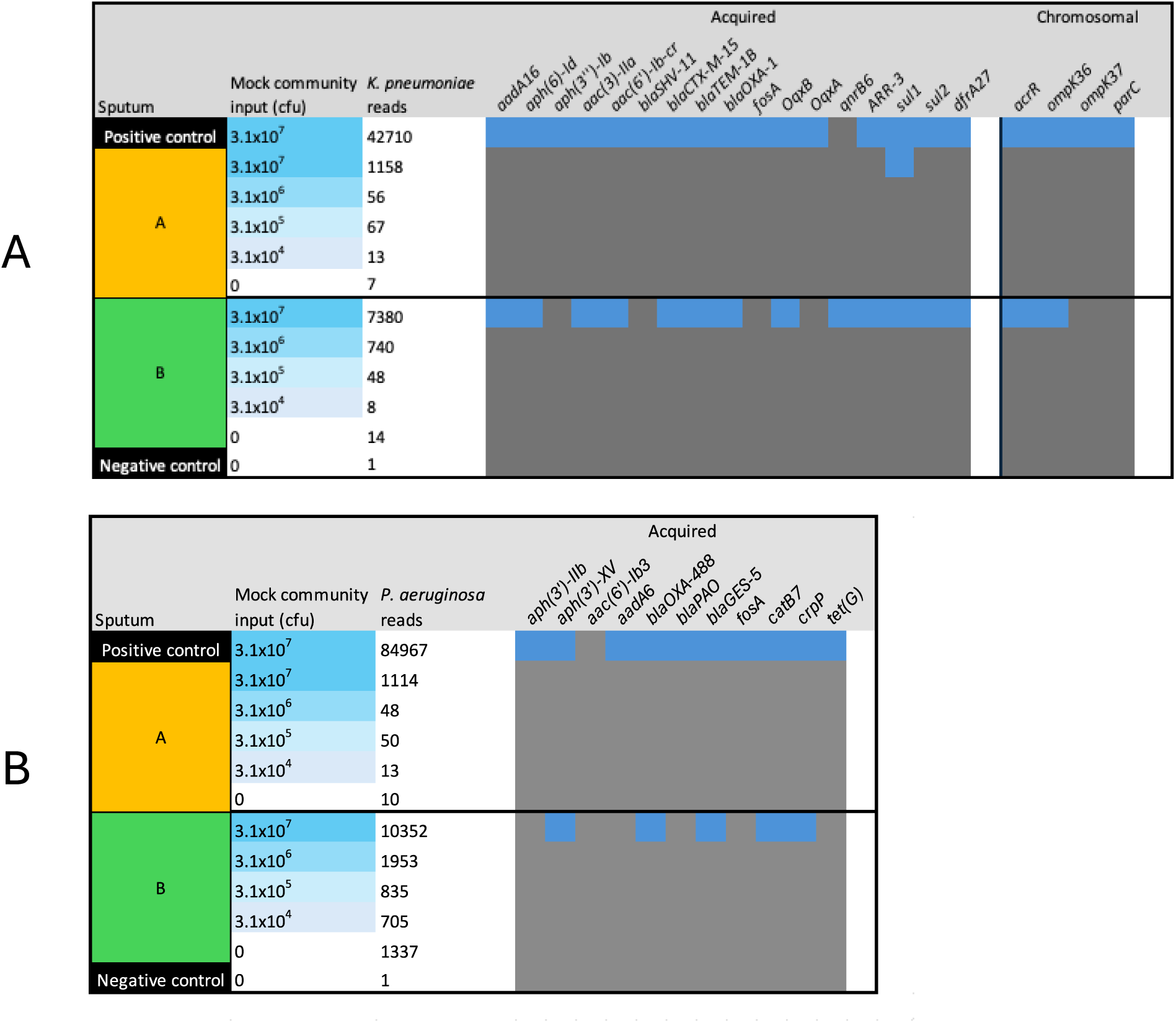
Detection of AMR gene profiles in two sputum samples. Genes in the header row represent AMR genes or AMR-linked chromosomal mutations previously detected in in-house isolate assemblies of **A**. *K. pneumoniae* and **B**. *Pseudomonas aeruginosa* used in the mock community

Sequencing of the negative control indicated two sources of contamination that need to be accounted for. *Lysobacter enzymogenes* accounted for 1405 reads, possibly originating from the achromopeptidase preparation used during DNA extraction, whereas 515 reads were taxonomically assigned to *Homo sapiens*, which could represent barcode leakage or trace cross-contamination from other high-host proportion samples multiplexed on the same flowcell (**Table S2**). Taxonomic classification of reads from the unspiked sputa also suggested that their innate bacterial loads were comparable to those of the spikes, with 33413 reads taxonomically assigned to *Staphylococcus haemolyticus* in sputum A, and >10^3^ reads assigned to species including *Enterococcus faecium, Lacticaseibacillus rhamnosus* and *Streptococcus dysgalactiae* in sputum A and *Streptococcus oralis, Streptococcus dysgalactiae* and *Pseudomonas aeruginosa* in sputum B.

In summary, we could detect >100 reads per million for each mock community species at a bacterial load of approximately 8×10^5^ cfu per species. Higher read counts of >10^4^ reads were required for comprehensive AMR profile detection, and we estimated that we would need >10^6^ total reads per sample assuming a pathogenic read abundance of ∼1%. As the total read counts per sample from the NAS half of the pilot flowcell ranged from 0.21-1.07×10^6^ (**Table S1**), we reasoned that this target read yield was attainable from multiplexing eight samples on one or two flowcells fully running NAS, assuming that the pilot run performance could be reproduced.

### Rapid pore loss greatly reduces sequencing output and metagenomic profiling performance

We next sequenced eight different clinical sputa (C1-C8)on two MinION flowcells running NAS on all channels to evaluate metagenomic profiling performance across samples of highly variable taxonomic composition. Contrary to our expectation that whole-flowcell NAS would improve read turnover and sequencing performance, we observed a reduction in sequencing output of ∼80% relative to the pilot sequencing run, obtaining 1.89M/2.19M reads and 2.2/2.8Gb output from both flowcells (**Table 1**). The pore activity reports indicated that sequencing pores were lost at an accelerated rate, with ∼90% of the initially available pores displaying the “inactive” status 24h into the run, and visual inspection of the flowcells indicated that this was not due to the introduction of air bubbles. Pores could be partially and temporarily rescued with the nuclease wash and sample reloading in flowcell 2B, but could only improve sequencing output by ∼25% relative to flowcell 2A (**Fig. S2**). This drastically reduced yield necessitated the pooling of sequencing output from both flowcells, reaching only 73.5% (5Gb) of the base output and 58.1% (4.18M) of the read count of the NAS half-flowcell in the pilot sequencing run.

Taxonomic composition varied greatly between samples, with total bacterial read counts ranging from 1.4×10^3^ – 4.2×10^5^ reads (median: 5.0×10^3^) and bacterial read proportion ranging from 0.27-85.8% (median:1.7%) (**Fig. 3A&B**). These distributions were heavily skewed by sample C5, which accounted for 87.2% of the bacterial reads sequenced across both flowcells (**Table 2**). Bacterial read count and proportion was generally higher in less viscous samples, which likely contained less host material (**Fig. 3B**). AMR genes were only detected in 2/8 of the samples, supporting our estimation from the pilot run that >10^5^ reads from the relevant species were needed for AMR detection (**Fig. 3C**, **Table 2**). Most of the isolate species identified from sputum culture could be detected from the samples at >100 reads (6/8 samples), and some species previously unidentified from the culture were detected at a similar level or higher. However, read counts of species identified from metagenomic sequencing generally fell short of our 10^5^ target required for comprehensive metagenomic profiling. Similar to the pilot run, negative control sequencing indicated read contamination or barcode cross-talk from host DNA and *Lysobacter enzymogenes*, and also *Serratia marcescens* which was highly abundant in C5 and *S. dysgalactiae*. These species would likely need to be subtracted from the output in actual clinical diagnostic and surveillance practice. In summary, rapid pore loss in the flowcells reduced total sequencing output by 5×, overriding the estimated 3× target enrichment from NAS and limiting metagenomic functionality to just species detection.

**Fig. 3.**
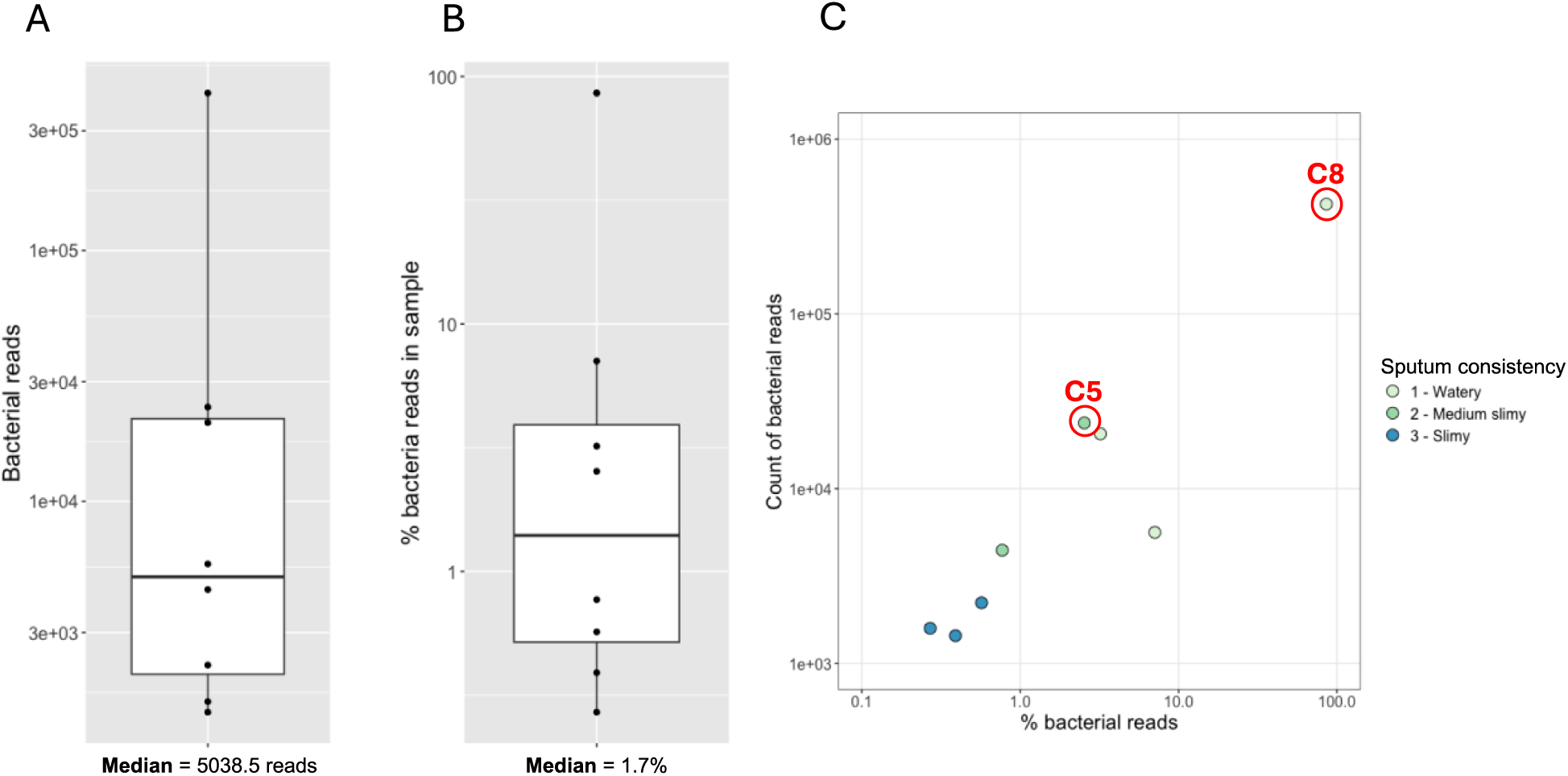
Variability in bacterial composition between different clinical sputa. **A** – Bacterial read count per sample. **B** – Proportion of bacterial reads per sample. **C** – Scatterplot summary of bacterial composition of eight clinical sputa. Sputa with detected AMR genes are circled in red.

**Table 2.**
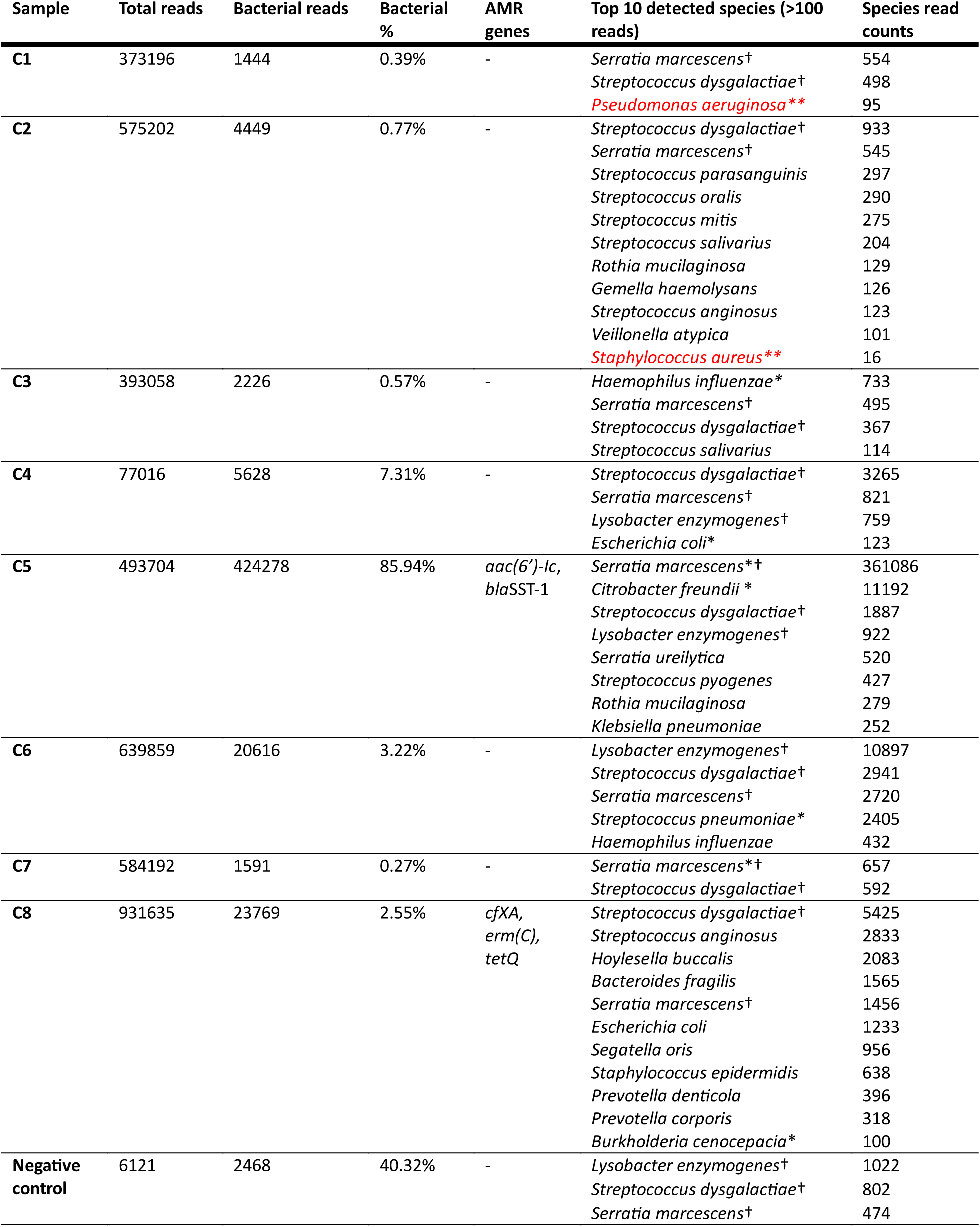
Taxonomic composition of NAS-sequenced clinical sputa. * - species identified from sputum culture, ** species identified from culture below detection threshold, † - likely contaminant species also detected in negative control

### Rapid pore loss cannot be mitigated by extensive sample decontamination measures

We consequently focused on troubleshooting pore loss in an attempt to improve metagenomic functionality. While rapid pore loss has frequently been anecdotally reported on the ONT community forums, especially in the context of the current generation R10.4.1 chemistry flowcells, its causes are still to-date poorly defined and documented in both the official ONT best practices guides and scientific literature. Consultation with the ONT technical support team suggested that **1.** pore loss could be accelerated by insufficient sample loading and incomplete pore occupancy during the run or **2.** caused by the presence of unknown contaminants from the sample. We also investigated **3.** any potential effects of NAS on flowcell health, due to previous reports(8, 9, 12) and the NAS best practices guide(10) indicating that NAS read rejection could result in reduced sequencing output, especially in the case of longer reads (>10kb).

We first addressed the likely sources of contamination from our samples. For this purpose, we used DNA extracted from four clinical sputa for our troubleshooting run, and switched from the earlier ligation sequencing kit with PCR-barcoding extension to a rapid barcoding kit protocol as it requires less input DNA per sample (1-4ng), allowing us to use a smaller volume of the extracted sputum DNA and hence reduce contaminant carryover from the sputum. We additionally increased the number of PCR cycles to 30× to further amplify sample DNA concentration relative to potential contaminants, and the pooled libraries were washed thrice using AMPure beads as recommended in the protocols. This resulted in a superficially high-purity DNA sample based on Nanodrop UV absorbance spectroscopy, with a 260/230 ratio of 1.92 and a 260/280 ratio of 1.93. In order to address point **1**, we loaded an increased amount of pooled library per flowcell (150fmol, 3× higher than the recommended 50fmol minimum) to maximize pore occupancy. The same library pool was loaded on four flowcells, with two flowcells running NAS on 50% of the channels to assess if NAS accelerated pore loss on the NAS half, and also two control flowcells without NAS to investigate there was accelerated global pore loss across the NAS flowcells relative to non-NAS setups.

Rapid pore loss was once again observed on both NAS and control flowcells despite all the troubleshooting measures taken, resulting in a 75% reduced sequencing output relative to the pilot run (**Fig. S3, Table 3**). As before, ∼90% of initially available pores were inactive 24h into the run and could not be adequately recovered by a nuclease wash/reload step. Pore occupancy was at ∼90% throughout the runs, suggesting that low pore occupancy was not the cause. In response to the poor performance of the first set of replicates, we treated the pooled libraries with RNAse A to rule out pore blockage by RNA molecules from the PCR reaction or the sample. This did not improve run performance in the second NAS cell, and while there was a modest increase in sequencing output in the second control flowcell (4.6Gb vs 3.3 Gb, 39%) it was difficult to tell whether this could also be due to inter-flowcell variability with a single replicate. Total sequencing output from the NAS flowcells was similar to that of the control flowcells, suggesting that NAS was not a major cause of pore loss in our situation.

**Table 3.**
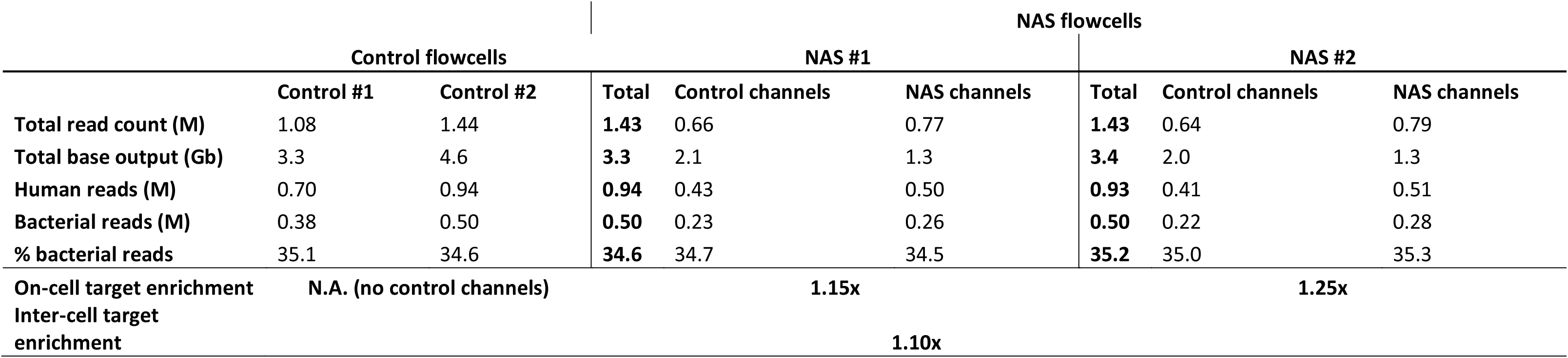
Summary statistics of troubleshooting runs.

NAS enrichment of bacterial reads was significantly lower than that of the pilot run, with an estimated 1.2× on-cell enrichment comparing NAS and control pores on the NAS flow cells, and a relative x enrichment in bacterial read yield in NAS flowcells relative to the control flowcells (Table 3). This limited enrichment was likely due one of the four clinical samples being 95% bacterial by composition, resulting in 35% target bacterial reads in the multiplexed library pool, which is higher than the recommendation of previous studies(8) and the ONT best practices guide of a 1-10% target composition to maximize potential enrichment gain. This highlights how a single deviant sample in a multiplexed setup can affect NAS enrichment across the entire flowcell. In summary, while we were able to rule out NAS and pore underloading as likely causes for the observed rapid pore loss, we could not eliminate the problem despite rigorous measures to reduce potential contaminants.

## Discussion

We evaluated the use of NAS for the enrichment of target bacterial genomic sequences from sputum metagenomic samples observing an enrichment of at best 3.1× by read yield, similar to previous metagenomic applications of NAS(9, 13–20). Whether this constitutes a functionally significant enrichment is dependent on the metagenomic problem at hand - under applications where the scientific question can already be answered without NAS, a three-fold enrichment introduced by NAS could indeed triple multiplexing capacity or improve the time-to-result in the context of rapid diagnosis. However, in our intended use-case of culture-free metagenomic surveillance of respiratory pathogens, this level of enrichment did not yield sufficient reads for the near-complete genomes required for AMR profiling in most of the clinical samples tested, due to the overwhelming proportion of contaminant host DNA of up to >99.7%. We suggest that NAS should thus be treated more as a low-cost polishing step in already-feasible workflows, rather than as a reliable replacement for wet-lab target enrichment protocols. Existing approaches for enriching low abundance respiratory pathogens currently outperform NAS – culture-based isolate sequencing remains the gold-standard in delivering pure high-coverage genomes, whereas established chemical sputum host-depletion protocols and target-capture using biotinylated baits yield superior target enrichment of up to 256× (4) and 300× (2) respectively, with the caveat of increased sample processing time and costs.

We multiplexed up to eight samples on the same flowcell to improve sample throughput and reduce overall sequencing costs, but in the process introduced two major complications to the workflow. Firstly, the presence of a few high bacterial load samples such as those in the troubleshooting run could raise the overall target composition on the flowcell above the 1-10% best-practices ideal and reduce the overall enrichment benefit across all samples loaded on the flowcell. Secondly, barcode cross-talk was observed in the sequence reads, with high-abundance species from one sample appearing at lower counts in the other samples and the negative control. This needs to be accounted for by manual curation, automatic bioinformatic methods or the use of dual barcoding systems with reduced barcode crosstalk. Given these limitations and the low target enrichment by NAS, we agree with the recommendation of a prior ONT respiratory metagenomics study to monoplex metagenomic samples for optimal data yield and performance(1), with the major caveat of increased sequencing costs.

We attempted to troubleshoot the rapid pore loss issue with the aim of improving metagenomic profiling with the potential 5-10× recovery in sequencing output, but were unable to resolve it or conclusively determine its cause. We ruled out pore underloading and NAS as probable causes, but could not prevent pore loss despite extensive sample decontamination measures. Two possible explanations remain, the first being that sputum samples might be inherently difficult to sequence with ONT, containing persistent inhibitors of pore activity that remain potent despite rigorous sample dilution and washing, and the second being that the undetermined pore inhibitor may have been introduced during library preparation. As the Nanodrop UV absorbance spectra of our pooled libraries conformed to what would be expected of a high-purity DNA sample, this unknown factor either has inconveniently similar absorbance characteristics to DNA, does not register on the absorbance spectrum, or is DNA itself. We speculate that sputum-origin inhibitors could include covalently-modified DNA, secondary DNA structures (e.g. hairpins), or sputum polysaccharides co-purified with the sample. PCR-based library preparation might also systematically introduce short barcoding primer-dimers that might affect flowcell performance. This could be exacerbated by the inclusion of a negative control and other low-concentration samples, which could promote primer-dimer formation due to insufficient PCR template DNA. Further research is needed to investigate if stringent DNA size selection or the omission of low-concentration samples can mitigate the problem.

## Conclusion

Rapid pore loss is a common cause of ONT run failure but remains undocumented in peer-reviewed literature. Our study indicates that rapid pore loss significantly reduces data output and limits metagenomic application, and that its idiosyncratic nature poses a challenge to our original goal of deployment in lower-middle-income countries, which often lack the necessary resources and logistical support to troubleshoot on-the-fly. We thus document our troubleshooting efforts to stimulate systematic discussion of the problem and further development, in the hope that long-read sequencing of clinical sputum samples will be attainable for surveillance and diagnostic applications in the future.

## Acknowledgements

This study was co-funded by the PPP allowance made available by Health∼Holland, Top Sector Life Sciences & Health (project number: LSHM22031) as part of the “AMR-Global: Reducing inappropriate exposure to antibiotics” (GLORIA-2) program. In addition, we acknowledge KNCV’s Dream Fund project “No more pandemics”, financially supported by the Dutch Postcode Lottery, for enabling KNCV’s contribution to this project. We also acknowledge the Utrecht Sequencing Facility (USEQ) for providing sequencing service and data. USEQ is subsidized by the University Medical Center Utrecht and The Netherlands X-omics Initiative (NWO project 184.034.019). Adri van der Zanden, founder and previous director of Mozand B.V., is very much thanked for his support to the start-up of this project.

## Supplementary Figures/Tables

**Fig. S1.**
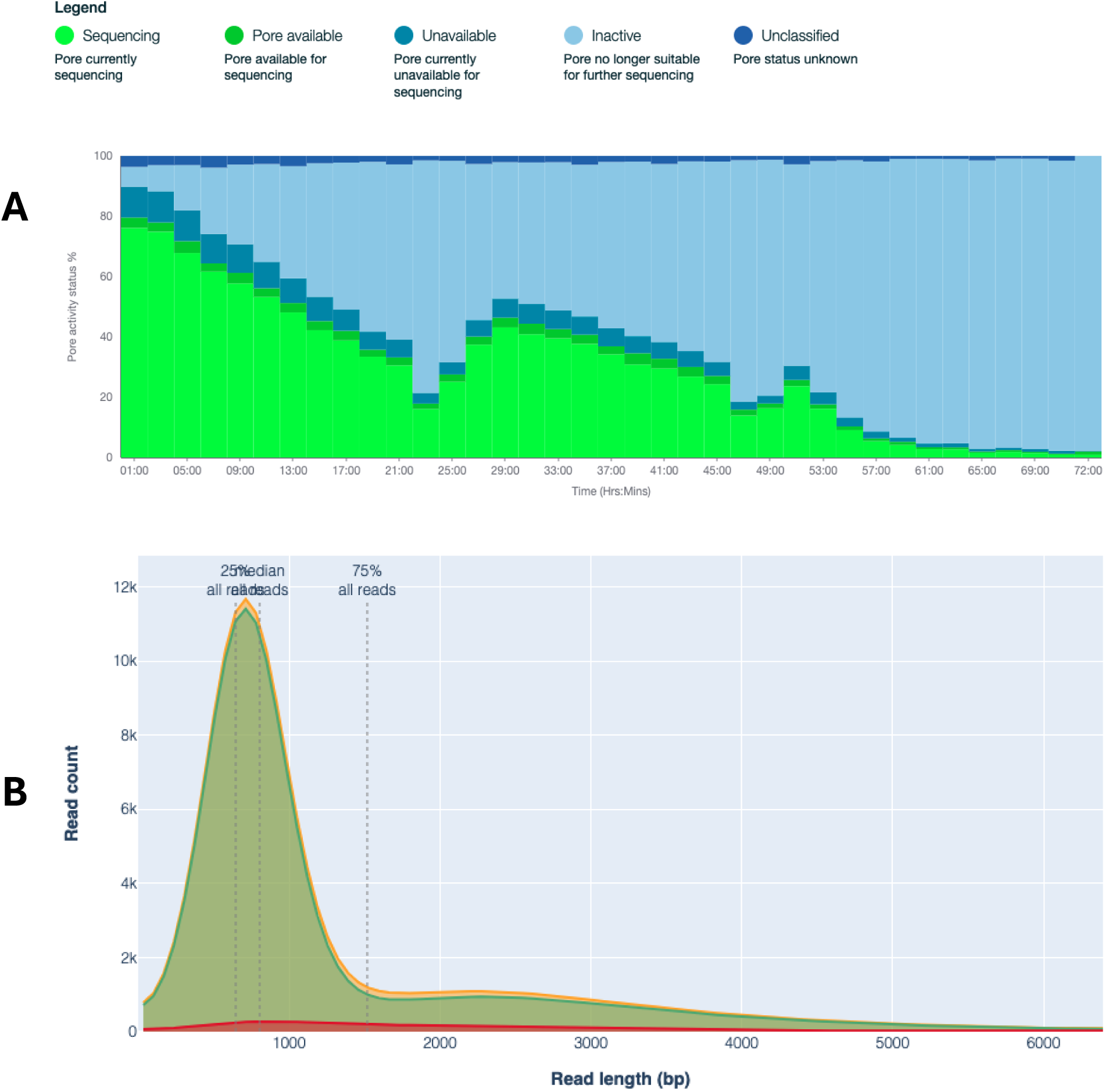
ONT sequencing reports for pilot run. **A.** Pore activity plot of pilot run flowcell. **B.** Bimodal read length distribution of NAS-sequenced reads with modes at ∼500bp (rejected reads) and ∼2500bp (kept reads)

**Fig. S2.**
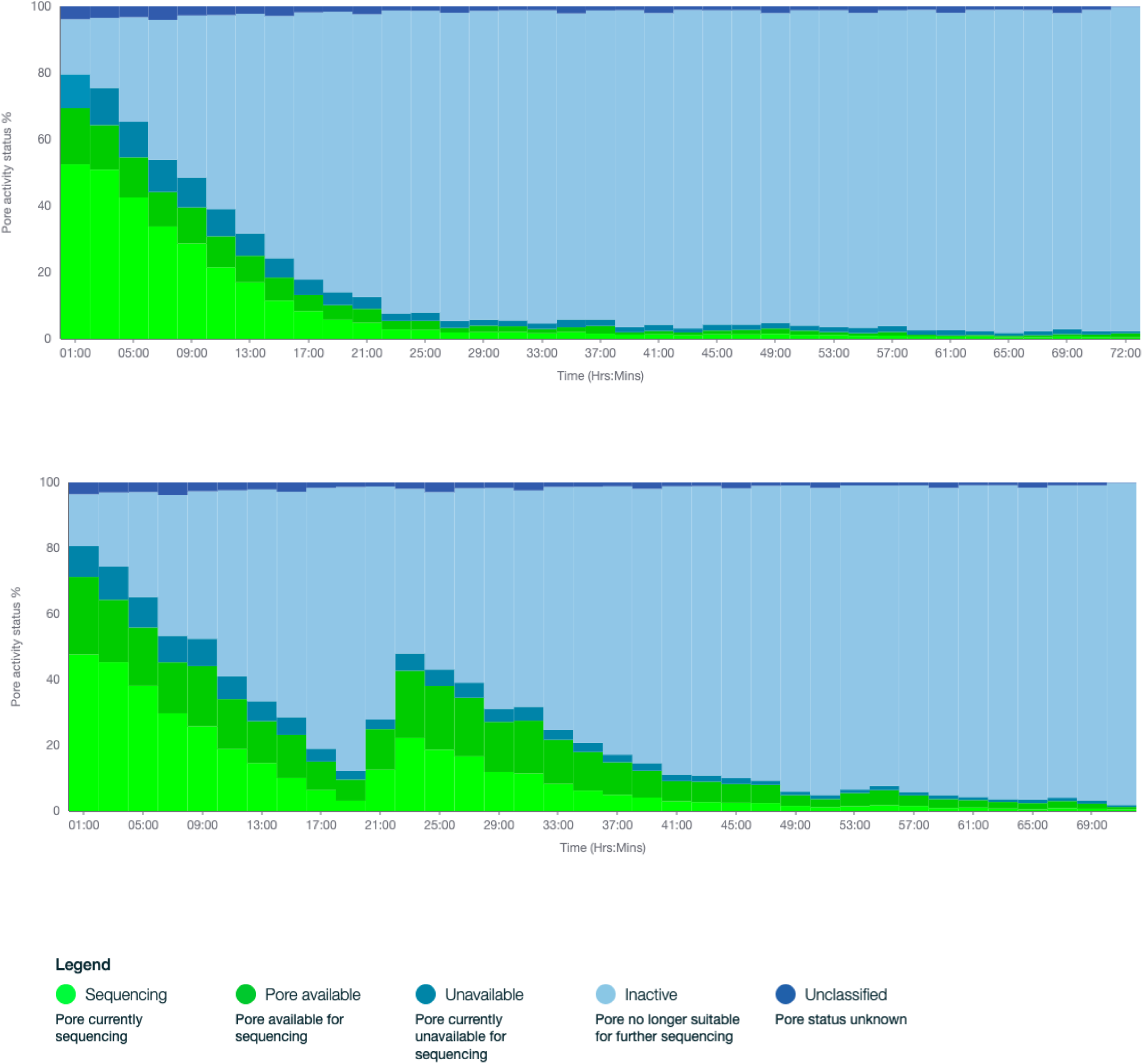
Rapid decline of pore activity in flowcells in second run.

**Fig. S3.**
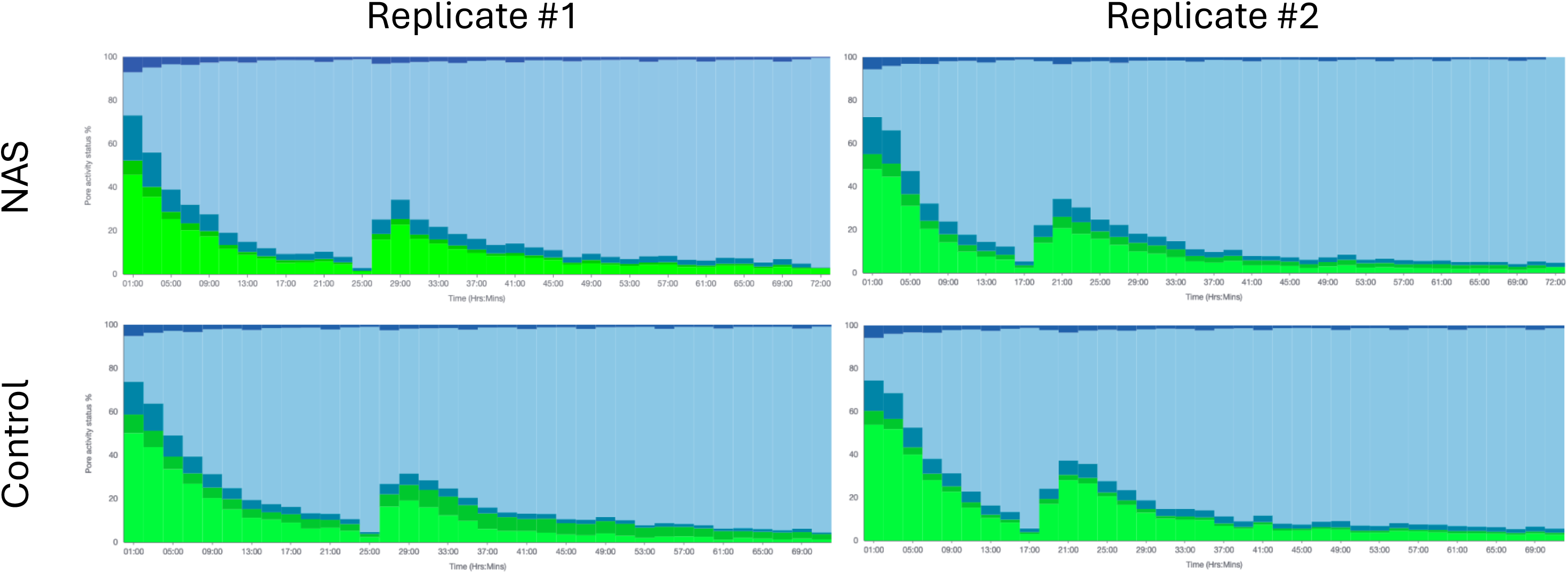
Pore activity plots for troubleshooting run flowcells.

**Table S1.** Per sample summary statistics of pilot NAS run. The MinION flowcell was split into control (1–256) and NAS channels (257–512)

| Sample | Sputum sample | Mock community spike (cfu) | Total |  |  | Control channels |  |  | NAS channels |  |  | Enrichment fold |  |
| --- | --- | --- | --- | --- | --- | --- | --- | --- | --- | --- | --- | --- | --- |
|  |  |  | Total read count | Bacterial read count | % bacterial reads | Total read count | Bacterial read count | % bacterial reads | Total read count | Bacterial read count | % bacterial reads | Bacterial read yield | Bacterial composition |
| **A1** | A | 3.10E+07 | 865490 | 38709 | 4.5 | 219875 | 9272 | 4.2 | 645615 | 29437 | 4.6 | 3.2 | 1.08 |
| **A2** | A | 3.10E+06 | 281855 | 16211 | 5.8 | 72705 | 4082 | 5.6 | 209150 | 12129 | 5.8 | 3.0 | 1.03 |
| **A3** | A | 3.10E+05 | 365556 | 9530 | 2.6 | 94109 | 2301 | 2.4 | 271447 | 7229 | 2.7 | 3.1 | 1.09 |
| **A4** | A | 3.10E+04 | 740609 | 49815 | 6.7 | 186975 | 12432 | 6.6 | 553634 | 37383 | 6.8 | 3.0 | 1.02 |
| **A5** | A | 0 | 1273359 | 64699 | 5.1 | 317931 | 15781 | 5.0 | 955428 | 48918 | 5.1 | 3.1 | 1.03 |
| **B1** | B | 3.10E+07 | 1433934 | 54488 | 3.8 | 362205 | 12896 | 3.6 | 1071729 | 41592 | 3.9 | 3.2 | 1.09 |
| **B2** | B | 3.10E+06 | 1344641 | 18578 | 1.4 | 347940 | 4467 | 1.3 | 996701 | 14111 | 1.4 | 3.2 | 1.10 |
| **B3** | B | 3.10E+05 | 738346 | 6255 | 0.8 | 185730 | 1426 | 0.8 | 552616 | 4829 | 0.9 | 3.4 | 1.14 |
| **B4** | B | 3.10E+04 | 1086484 | 6281 | 0.6 | 270277 | 1452 | 0.5 | 816207 | 4829 | 0.6 | 3.3 | 1.10 |
| **B5** | B | 0 | 1240193 | 9384 | 0.8 | 312800 | 2147 | 0.7 | 927393 | 7237 | 0.8 | 3.4 | 1.14 |
| **Positive control** | NA | 3.10E+07 | 244945 | 243286 | 99.3 | 59133 | 58710 | 99.3 | 185812 | 184576 | 99.3 | 3.1 | 1.00 |
| **Negative control** | NA | 0 | 2647 | 1984 | 75.0 | 621 | 473 | 76.2 | 2026 | 1511 | 74.6 | 3.2 | 0.98 |
| **Total** |  |  | 9618059 | 519220 | 5.4 | 2430301 | 125439 | 5.2 | 7187758 | 393781 | 5.5 | 3.1 | 1.06 |

**Table S2.** NAS decision performance statistics. Proportions of correct/incorrect NAS decisions are indicated on the confusion matrix.

|  | Human | Bacteria |
| --- | --- | --- |
| Reject | True positive<br>$6.44 \times 10^6$<br>(89.7%) | False positive<br>$2.03 \times 10^4$<br>(0.3%) |
| Keep | False negative<br>$3.47 \times 10^5$<br>(4.8%) | True negative<br>$3.73 \times 10^5$<br>(5.2%) |

**Accuracy** (% correct calls made): **94.9%**
**Recall** (% human reads that are rejected): **94.9%**
**False positive rate** (% bacteria reads rejected): **5.2%**
**Negative predictive value** (% kept reads that are bacterial): **52%**

**Table S3.**
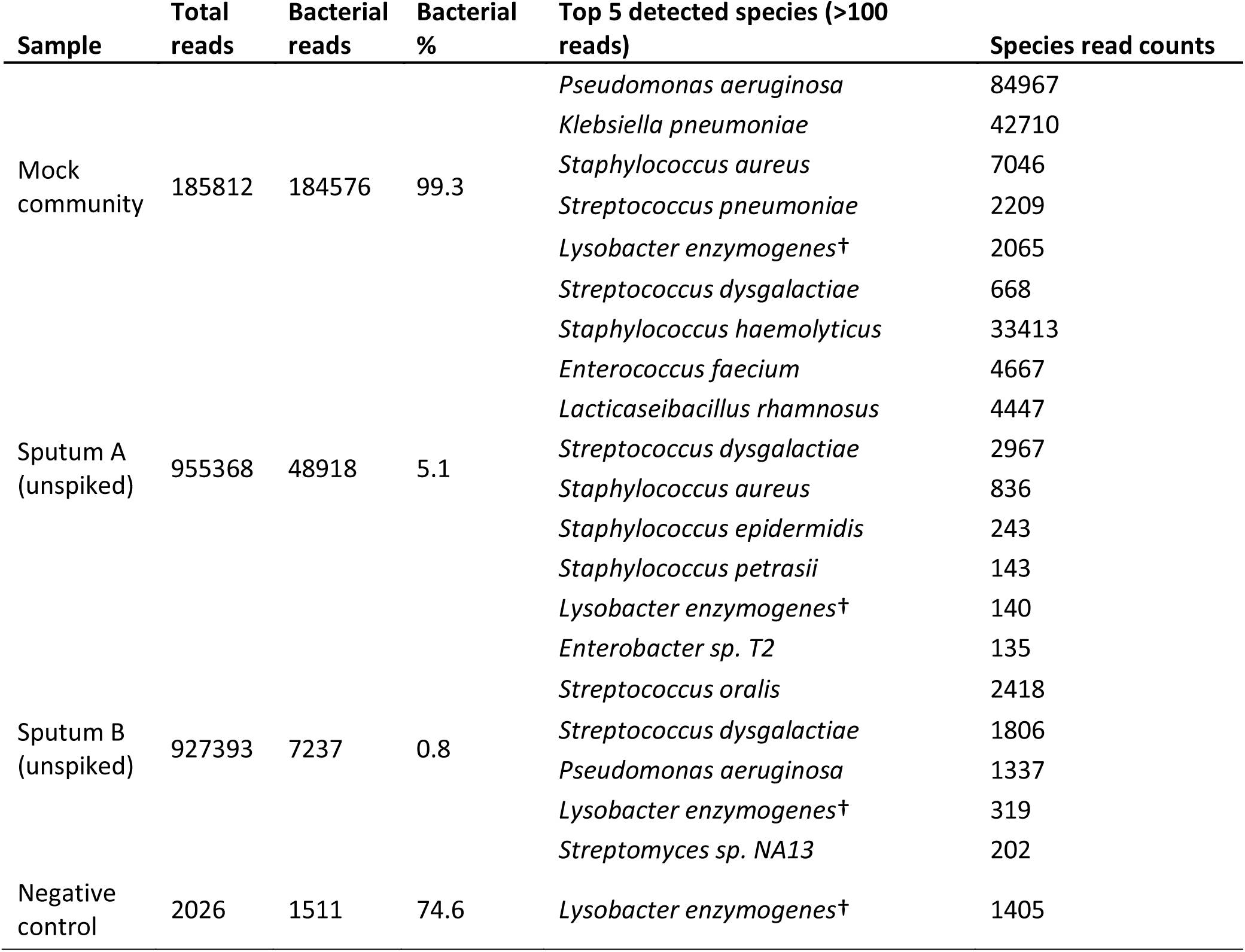
Taxonomic composition of pilot control samples. † - likely contaminant species also detected in negative control.

## Notes

### Competing Interest Statement

The authors have declared no competing interest.

